# Temporal and spatial dynamics mapping reveals follicle development regulated by different stromal cell populations

**DOI:** 10.1101/2022.03.04.480328

**Authors:** Xiaoqiang Sheng, Jidong Zhou, Nannan Kang, Wenwen Liu, Lina Yu, Zhe Zhang, Yang Zhang, Qiuling Yue, Qiwen Yang, Xinke Zhang, Chaojun Li, Guijun Yan, Haixiang Sun

## Abstract

Follicle development is a complex dynamic process. The follicles are encapsulated in the stroma, and once the follicle develops, the follicle moves from the cortex to the medulla and finally to the cortex to ovulate in the process of continuous growth. Many of these processes cannot be explained with the follicle alone. Through single-cell and spatial transcriptome sequencing at key time points of follicular development in mice after birth, we found that ovarian stromal cells are not only one of the main cell groups that make up the ovary but that their cell population and spatial location are also closely related to follicular development. Through analysis of cell communication, it was found that ovarian stromal cells were the main transmitters of intercellular communication, and many of the signals they sent were received by granulosa cells and oocytes to participate in follicle development. Ovarian stromal cells are not a homogeneous cell population. We combined single cell types with their spatial location information to divide ovarian stromal cells into four types, namely, structural stromal cells, perifollicular stromal cells, stromal progenitor cells, and steroidogenic stromal cells, each of which plays a different function in follicle development. Indepth studies of the different spatial locations and different types of stromal cells will expand our understanding of follicle development dynamics, leading to new targets and novel approaches for the treatment of ovarian-related diseases.

## Introduction

The ovary comprises two components: the parenchyma and the stroma. The follicle is the functional unit of the ovary and comprises the ovarian parenchyma, which determines the reproductive lifespan of females^1^. It is composed of an oocyte and layers of somatic cells (granulosa and theca cells) embedded in the ovarian stroma. The bulk of the ovary, both the cortex and medulla, consists of stroma with different characteristics. Most follicles are embedded in the stroma of the ovarian cortex and mature in the stroma of the ovarian medulla^2^. Historically, research in ovarian biology has mainly focused on folliculogenesis controlled by oocytes and granulosa cells (GCs), while there are many unsolved problems in folliculogenesis, which cannot be explained by only these two cell types. In addition to cells making up the follicle, the ovarian stroma in which the follicles are embedded contributes greatly to folliculogenesis but does not receive enough attention, as it holds critical keys to understanding the complex process of folliculogenesis.

The ovarian stroma is composed of general components, such as immune cells, blood vessels, nerves, extracellular matrix, and a mixed population of incompletely characterized stromal cells^3^. The majority of the ovarian stroma comprises incompletely characterized stromal cells that include fibroblast-like, spindle-shaped, and interstitial cells^4^. Ovarian stromal cells have significant roles in folliculogenesis, particularly in the activation of primordial follicles and the differentiation of theca cells^5, 6^. Secondary follicles can be grown alone using in vitro culture systems, while primordial and primary follicles must be activated in organ cultures containing stromal components^7^. Moreover, all studies that successfully resulted in the production of human metaphase II (MII) eggs following in vitro culture used mechanical methods rather than enzymatic digestion to isolate follicles from the ovarian tissue, because enzymatic digestion causes damage to the follicles while also maintaining the close apposition of residual stromal cells, which engage in intensive intercellular communication with the follicle and may have the capacity to differentiate into other critical cell types, such as theca cells^8^. Furthermore, stromal cell coculture can improve the growth and survival of early follicles, which is critical for the successful translation of in vitro culture systems into primate and human follicles. Ovarian stromal cells not only provide structural support for follicle development but also have complex bidirectional paracrine signaling with the follicle^9^. Several growth factors are key regulatory molecules in folliculogenesis, including fibroblast growth factor, transforming growth factor beta, platelet-derived growth factor, hepatocyte growth factor, and insulin-like growth factor^10, 11^. Nevertheless, the specific cell–cell secreted signaling between follicle and stromal cells remains unclear.

Despite these new insights, our comprehension of ovarian stromal cells is still limited, and many questions remain unanswered. The incomplete characterization and categorization of ovarian stromal cells leads to confusion across studies that report findings about stromal cells without further identification. Ovarian stromal cells do not refer to a single homogenous cell population^3^. Recent single-cell RNA-sequencing (scRNA-seq) studies confirm the presence of multiple stromal cell clusters, while a comprehensive and complete characterization of stromal cell types throughout the ovary is lacking. The distribution and subtypes of stromal cells will likely differ with their location in the ovary (e.g., cortex vs. medulla). The stromal cell distribution is also likely to change, accompanied by cyclic structural changes as follicles grow and ovulate and the corpus luteam develops^12^. Changes are also evident over the whole reproductive lifespan, including increases in fibrotic collagen, as demonstrated in aged mouse, women, and primate ovaries^13–15^. For these incompletely characterized stromal cell types, careful ontology, further marker identification, and attention to nuances of regionality are critical next steps.

Although scRNA-seq can obtain information on the “temporal dynamic expression” of genes at the single-cell level, it still inevitably loses the spatial information of tissue samples. Thus, we cannot obtain enough information on stromal cells since they exhibit spatial variation during follicle development. With developments in spatial transcriptomics technologies, we have the ability to better characterize ovarian stromal cells and changes during follicular development and can refer to them with more precise names as we understand their individual roles in physiological and pathological processes. Thus, we conducted scRNA-seq and spatial transcriptomics analyses covering five important time courses that represent different follicle development stages to fully understand the role of ovarian stromal cells in follicle development and to completely characterize and categorize ovarian stromal cells.

In this research, we generated a single-cell map and spatiotemporal gene expression dynamics of the follicle development of the postnatal ovary in mice and elucidated the important role of ovarian stromal cells in follicle development. Cell–cell communication analysis delineates the complex intercellular communication relationships between groups of cell types at different stages of follicle development and reveals that ovarian stromal cells are the main outgoing signaling cell type in the ovary that regulates the development of follicles. We completely characterized the ovarian stromal cells into four subpopulations, namely, structural stromal cells, perifollicular stromal cells, stromal progenitor cells, and steroidogenic stromal cells, with further marker identification. Each of these stromal cells subpopulations play different role in follicle development.

## Result

### A single-cell map and spatiotemporal gene expression dynamics of follicle development in the postnatal ovaries of mice

To generate a cellular map of the mouse ovary that accounts for the temporal and spatial changes across follicle development, ovary samples were analyzed by single-cell RNA-sequencing (scRNA-seq) and spatial transcriptomics methods (10× Genomics Visium slides and high-resolution microscopy) (Figure 1 A). Ovaries at five time points were integrated into our scRNA-seq analysis: postnatal day (day 3), day 5, day 7, 3 weeks, and 8 weeks. Ovaries at three time points were integrated into our 10× Visium analysis: postnatal day (day 7), 3 weeks, and 8 weeks. The latter approach allows exploration of cell signatures in follicle development to obtain detailed cell populations and temporal changes.

**Figure 1.**
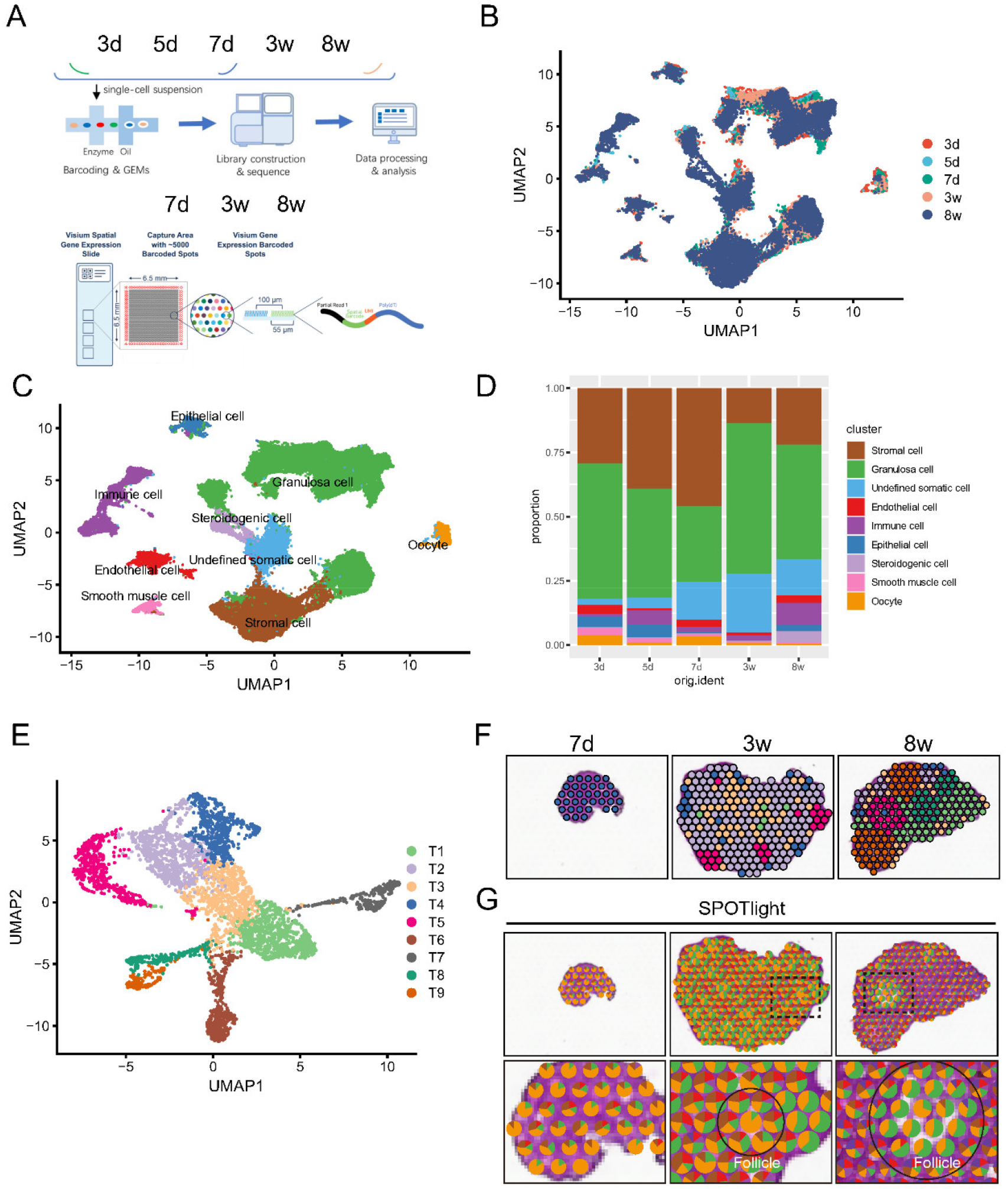
A single-cell map of follicle development in the postnatal ovaries of mice (A) Temporal and spatial dynamics mapping strategy of the mouse ovary. Ovaries were isolated and disaggregated into single-cell suspensions; cells were barcoded and used for library construction onto the 10× Genomics platform, and the data produced after sequencing were analyzed by dedicated software; (B) UMAP plot of ovarian cells colored by the selected developmental time points (stage); (C) UMAP plot of 9 ovarian cell types. (D) Percentages of the 9 ovarian cell types at day 3, day 5, day 7, week 3 and week 8. (E-F) UMAP plot and SpatialFeaturePlot of ovarian spots colored by cluster at 10× magnification. (G) Spot picture of the ovary in 7-day, 3-week, and 8-week sections.

We were able to generate an ovary map of 58,319 cells and to identify twenty-one clusters that were assigned cell identity based on their expression of markers. To further identify the clusters, we assigned the clusters based on known cell-type marker genes (Figure S1A). The cell-specific marker genes were oocyte (Sycp3), granulosa cell (Amhr2), stromal cell (Col1a1), endothelial cell (Cdh5), epithelial cell (Krt19), immune cell (Ptprc), smooth muscle cell (Des), and steroidogenic cell (Ptgfr). These clusters can be grouped into nine main cellular categories: (i) stromal cells, (ii) granulosa cells, (iii) oocytes, (iv) endothelial cells, (v) epithelial cells, (vi) immune cells, (vii) smooth muscle cells, (viii) steroidogenic cells, and (ix) undefined somatic cells (Figure 1C). To further obtain the dynamic pattern of the gene signature, the nine main cellular category cluster-specific marker genes were identified (Table S1). Among these cell types, stromal cells and GCs accounted for the vast majority of the proportion (Figure 1D).

To systematically map the location of the cell types identified by scRNA-seq within the ovary, we used 10× Visium Spatial Transcriptomics technology. We examined three ovary samples on day 7, week 3, and week 8, which represent three important ovarian development stages. Overall, nine transcriptionally distinct clusters were classified and visualized using Uniform Manifold Approximation and Projection (UMAP, Figure 1E), and their locations in the ovary are visualized in Figure 1F. We can clearly see the location of the nine clusters, accompanied by the follicle and corpus luteum. Then, we integrated spatiotemporal data with scRNA-seq data to infer the locations of cell types and states within the complex ovary tissue using SPOTlight. As shown in Figure 1G, we mapped the eight clearly identified cellular categories into the 10× Visium spatiotemporal ovary tissue and found that stromal cells, endothelial cells and GCs were the three cellular categories around the oocyte. We can see all the spatial locations of the ovarian cell types. As a resource, we can provide enough spatiotemporal information on various cell types for researchers in this field.

### Genetic dynamics signatures of the granulosa cell lineage during follicle development

Granulosa cells (GCs) are an indispensable part of follicles that directly interact with oocytes. To dissect the heterogeneity of GCs during development in the period of investigation through UMAP analysis, the GC populations were identified as 11 clusters (Figure 2A and B). To obtain the dynamic pattern of the gene signature, the GC cluster-specific marker genes were identified (Table S2). The top 10 expressed genes of each cluster are shown in a heatmap in Figure S2A. According to these analyses, a genetic dynamic model, including less differentiated, mural, cumulus, steroidogenic and proliferative GCs during follicle development, was drawn (Figure 2C). Genes known to be expressed in early development stage GCs are Wt1 and Foxl2; for mural GCs, Cyp19a1; for cumulus cells, Slc38a3 and Amh; and for steroidogenic GCs, Cyp11a1 and Lhcgr (Figure S2B). The marker genes Wt1, Slc38a3, Cyp19a1 and Cyp11a1 were visualized in 10× Visium sections, which matched the real histological location (Figure 2D). The proportion of less differentiated GCs decreased over time, and the proportions of cumulus GCs, mural GCs, and steroidogenic GCs increased over time and peaked at 3 weeks or 8 weeks (Figure 2E). The results matched the GC dynamics during ovary and follicle development. However, we did not find any distinct boundary to group the GCs across primordial to antral follicles.

**Figure 2.**
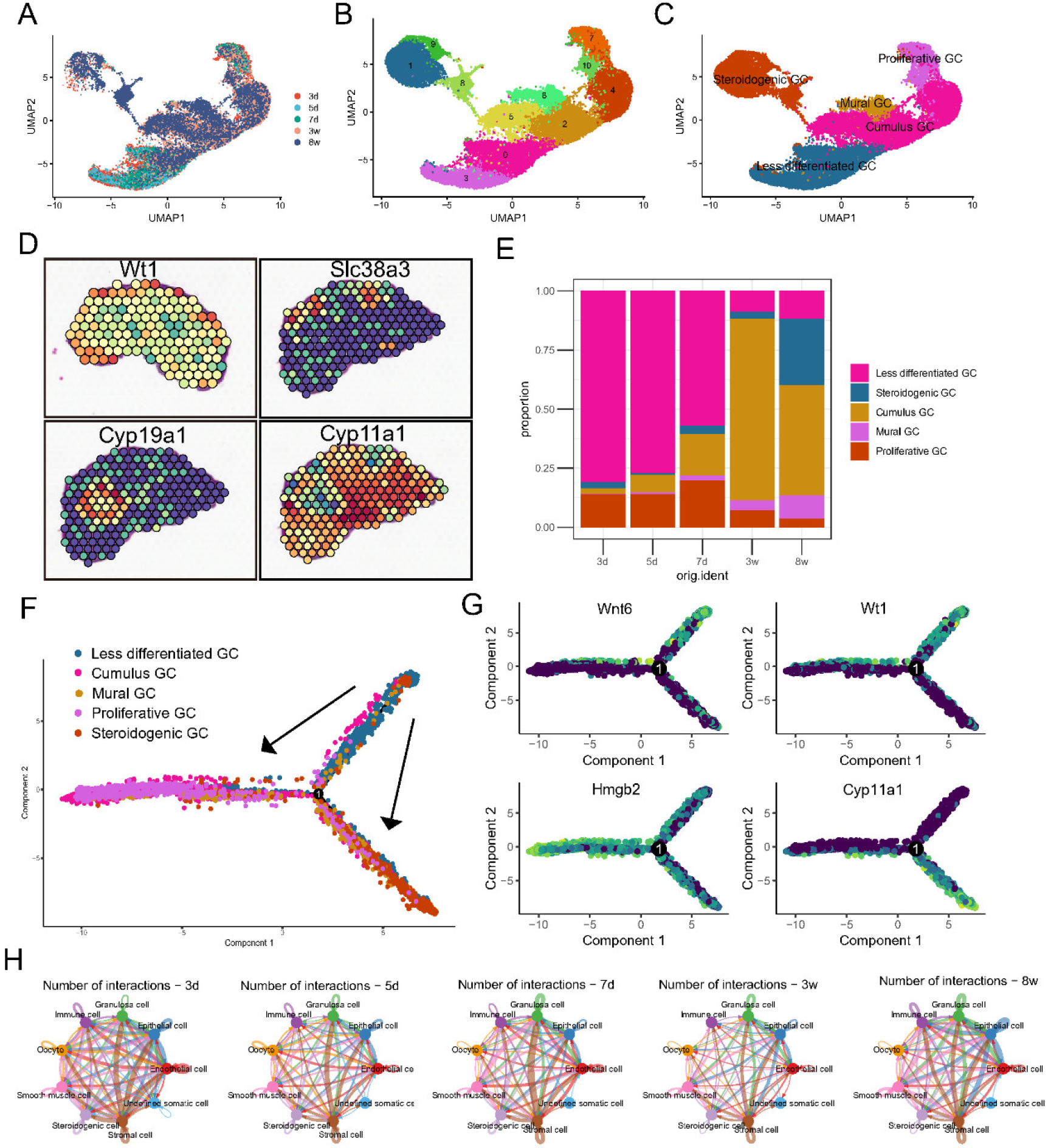
Genetic dynamics signatures of the granulosa cell lineage during follicle development (A) UMAP plot of GCs colored by the selected developmental time points (that is, stage); (B) UMAP plot of GCs colored by cluster; (C) UMAP plot of 5 GC types; (D) The location of Wt1, Slc38a3, Cyp19a1 and Cyp11a1 in 10× Visum ovary sections; (E) Percentages of the 5 ovarian cell types at day 3, day 5, day 7, week 3 and week 8. (F) Single-cell trajectories of the GC subsets as a function of developmental timeline and cell state. (G) Expression of developmental marker genes along pseudotime trajectories. (H) The number of interactions of the 9 ovarian cell types during the time course of 3 days, 5 days, 7 days, 3 weeks, and 8 weeks.

To dissect the fate determination of GCs throughout the investigated period, they were ordered along pseudotime trajectories according to the gene expression reported above. Three states (a term used in pseudotime analysis, where “state” is assigned to mark the segment of the trajectory tree in Monocle) were obtained. Moreover, the expression of representative genes (marker genes of the 3 follicle formation stages identified above) along with the pseudotime trajectories was consistent with our inference. The root state contained most of the less differentiated GCs; state 1 contained most of the cumulus GCs, mural GCs, and proliferative GCs; and state 3 contained most of the steroidogenic GCs (Figure 3F; Figure S2C). In such pseudotime trajectories, 3 branches implied 3 differentiated stages of GCs with cell markers, such as Wnt6, Wt1, Hmgb2, and Cyp11a1 (Figure 2G). GO analysis of representative genes of the three states showed the associated pathways (Figure S2D-F). Furthermore, we analyzed the cell crosstalk between the two major cell populations (stromal cells and GCs) and other cell types. As shown in Figure 2H, in addition to extensive intercellular communication between GCs and other cell types, there was also much stronger intercellular communication between stromal cells and other ovarian cell types.

**Figure 3.**
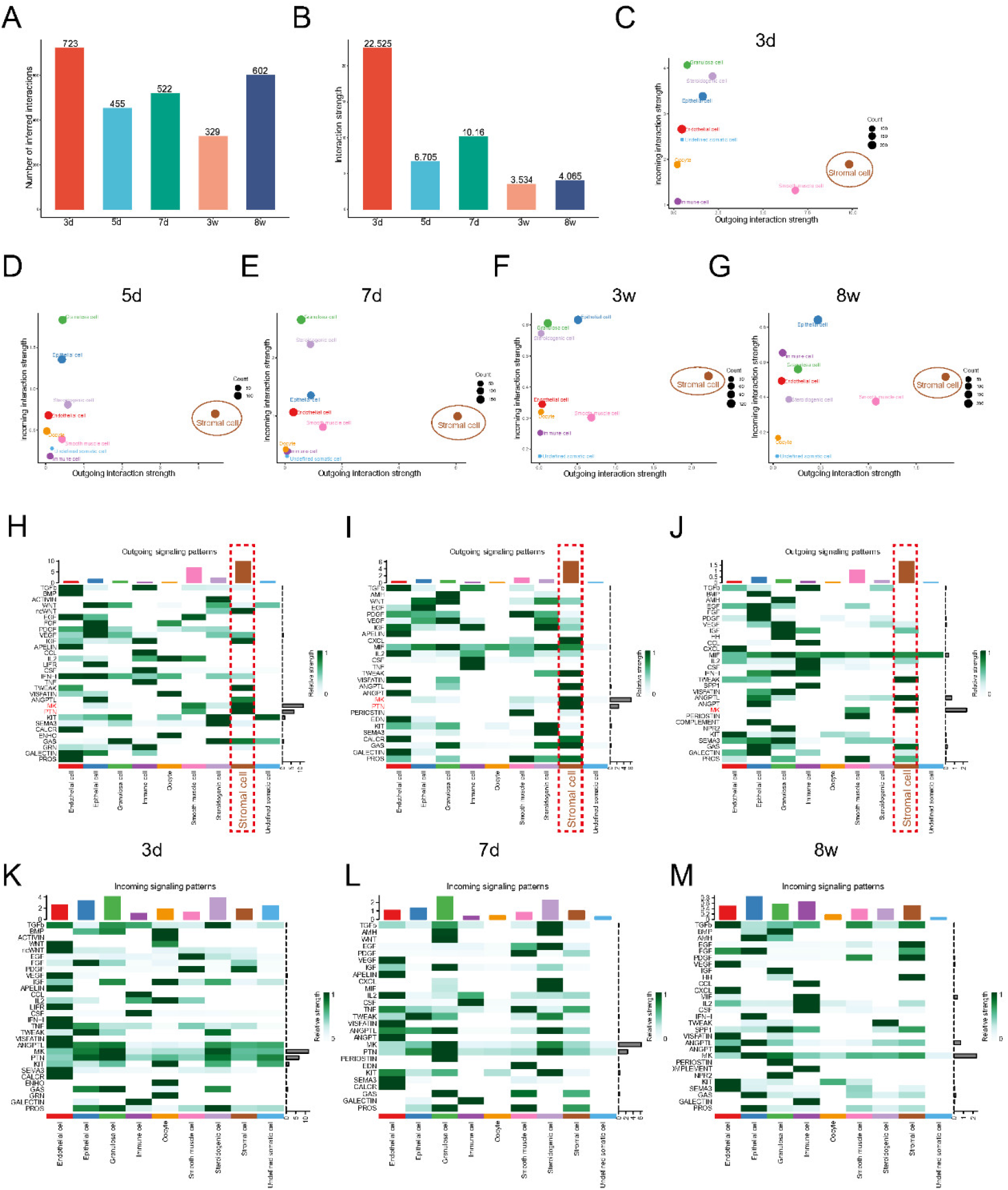
Ovarian stromal cells are the main outgoing signaling cell type in ovarian cell communication during follicle development (A and B) The number of inferred interactions and the interaction strength at day 3, day 5, day 7, week 3 and week 8. (C-G) The circle plot of the number of interactions among the 9 ovarian cell types at day 3, day 5, day 7, week 3 and week 8. (H-L) Signaling role analysis of the 9 ovarian cell types at day 3, day 5, day 7, week 3 and week 8. (M-O) The outgoing signaling patterns of the 9 ovarian cell types at day 3, day 5, day 7, week 3 and week 8.

### Ovarian stromal cells are the main outgoing signaling cell type in ovarian cell communication during follicle development

To analyze the communication of these cell types in the mouse ovary, we further performed CellChat analysis to identify the cell communication among these cell types during follicle development. As shown in Figure 3A-B, the number of inferred interactions on day 3, day 5, day 7, week 3, and week 8 was 723, 455, 522, 329, and 602, respectively, and the interaction strengths on day 3, day 5, day 7, week 3, and week 8 were 22.525, 6.705, 10.16, 3.534, and 4.065, respectively. We found that day 3, day 7, and week 8 were the strongest time points for cell communication. Thus, we focused more on the day 3, day 7, and week 8. There was very complex cell communication between the nine cell types in the ovary, and stromal cells were the cell type with the strongest signaling. Further signaling role analysis revealed that stromal cells exhibited the strongest outgoing interaction in cell–cell communication throughout the six important ovary development times (Figure 3C-G). While GCs were the cell type that received the most incoming receptor signals on days 3, 5 and 7, and epithelial cells were the type that received the most incoming receptor signals at 3 weeks and 8 weeks. Furthermore, we explored specific types of cellular communication across the selected time course. It was found that stromal cells frequently communicate with all other cells. In particular, the PTN, MK, ncWNT, PROS, GAS, ANGPTL, and TWEAK signaling pathways were actively involved in outgoing signaling patterns of stromal cells (Figure 3H-M; Figure S3). Among the outgoing signaling, MK is expressed during the entire follicle development stage. PTN was only detected on day 3, day 5, and day 7, when primordial follicle activation and follicle development to primary and secondary follicles were initiated (Figure 3H-M; Figure S3). As shown in Figure S4, we compared day 3 with day 5, when primordial follicles were activated, and found that ncWnt, PROS, APELIN, GRN, CALCR, ACTIVIN, IFN-1, and LIFR were specifically expressed on day 3, while IL-1 and ANNEXIN were specifically expressed on day 5. Other signaling pathways, such as TWEAK, PTN and MK, were expressed at higher levels than on day 5. When comparing the relative information flow of day 7 with that of day 5, we found that IL-1, ANNEXIN, CCL, BMP, FGF, and ENHO were specifically expressed on day 5, while TWEAK, APELIN, ANGPT, CALCR, CXCL, AMH, PERIOSTIN, PROS, EDN, and MIF were specifically expressed on day 7. Other signaling pathways, such as PTN and MK, were expressed at higher levels on day 7 than on day 5. We also compared the signaling on day 7 with that at 3 weeks and revealed that PTN, TWEAK, EDN, MIF, TNF, WNT, PERIOSTIN, CXCL, CALCR, APELIN, IL-2, and CSF were specifically expressed on day 7, while NPR2 and HH were specifically expressed on week 3. These cell signaling pathways changed during specific follicle development stages, suggesting that the secretory signals may contribute to the follicle development stage. Secretory factors like MK, secreted by stromal cells persist along with follicle activation to full follicle growth, which may represent the vital communication sent by stromal cells to support follicle development. We had mapped the crucial signaling pathways in follicle development. Thus, the cell communication among these cell populations during follicle development provides more information about the regulatory network of follicle development.

### Identification and genetic dynamics of ovarian stromal cell subpopulations

Furthermore, we analyzed the genetic dynamics of the ovarian stromal cell. When the transcriptome of the stromal cell was plotted along with the developmental stages, 11 cell clusters were identified (Figure 4A-C). As noted, novel marker genes within each cluster were identified (Table S3), whose expression varied according to developmental stage (Figure 4D). CytoTRACE analysis showed that clusters 6, 7, and 8 may be the root of stromal cells (Figure 4E). As shown in Figure 5F, feature plot analysis of the special cell markers of the identified clusters showed that Tcf21 expression was highest across the ovarian stroma, and Lum expression was mainly in clusters 0, 1, 2, 4, 5, and 10. The expression of Star and Cyp17a1 was mainly in clusters 1 and 9. The expression of Enpep was mainly in clusters 3, 7 and 8. The expression of Aldh1a2, a stem cell marker, was mainly in cluster 6. The genes Lum, Cyp17a1, Enpep and Aldh1a2 were visualized in 10× Visium sections and verified using in situ hybridization (ISH), immunohistochemistry (IHC) or immunofluorescence (IF) (Figure 4G-J). Lum expression in the stromal cells around the follicles supports the growth of the follicle. Cyp17a1 was located not only in theca cells around the follicle but also in some stromal cells outside the theca. Additionally, Enpep expression was associated with the development of follicles and was confirmed by ISH. The expression of Aldh1a2 was mainly in the ovary epithelium and the stroma of the ovarian cortex. The results suggested that ovarian stromal cells do not refer to a single homogenous cell population and can be clearly classified with the help of scRNA-seq and spatial transcriptomics methods.

**Figure 4.**
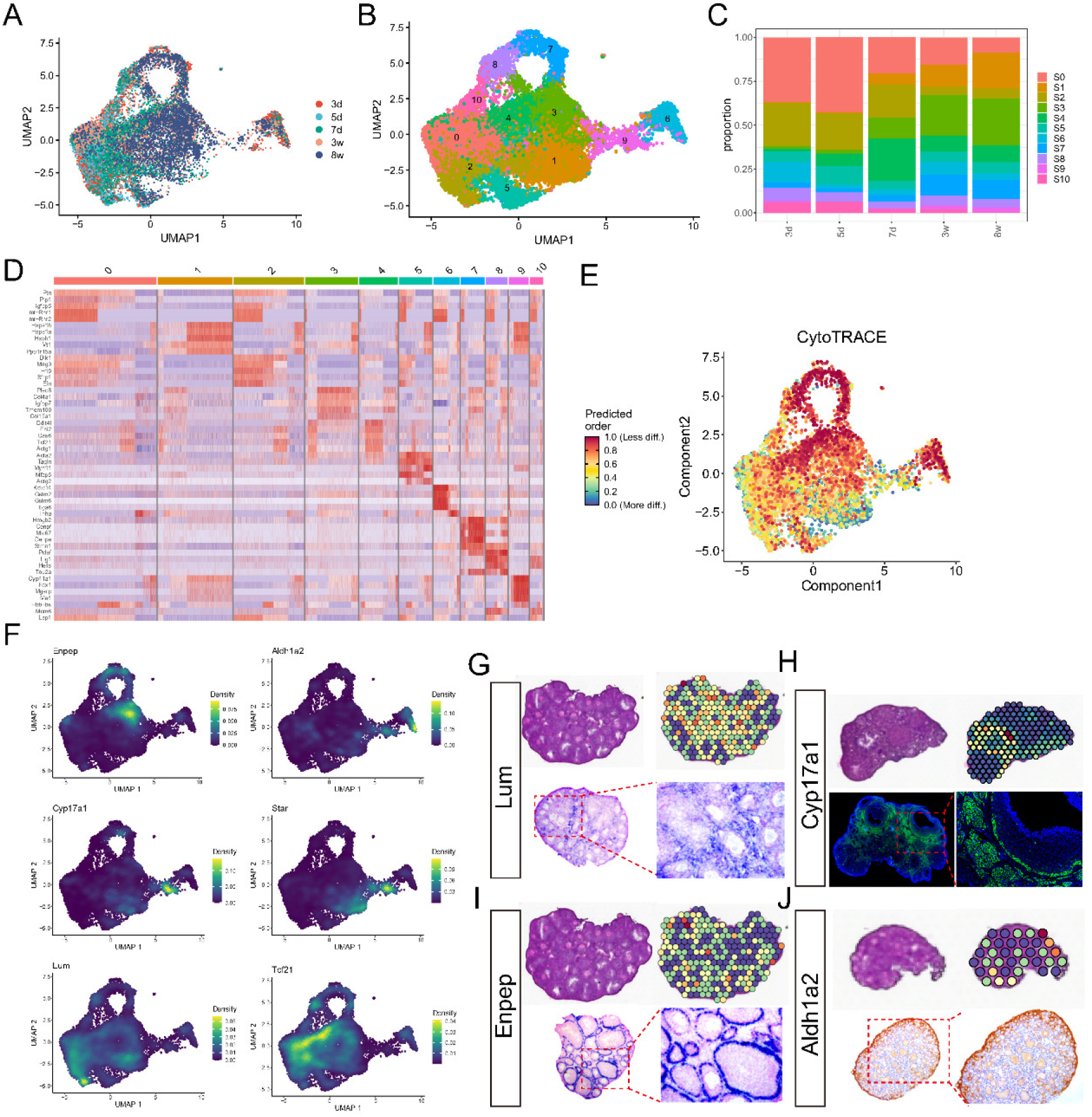
Genetic dynamics of ovarian stromal cell subpopulations (A) UMAP plot of ovarian stromal cells colored by the selected developmental time points; (B) UMAP plot of GCs colored by cluster; (C) Percentages of the 11 ovarian stromal cell clusters at day 3, day 5, day 7, week 3 and week 8. (D) Heatmap of the top 5 marker genes of the stromal cell clusters. The top 10 marker genes in each cluster are shown in Table S3. (E) CytoTRACE analysis for reconstructing cellular differentiation trajectories of the 11 clusters. (F) Feature plots of specific marker genes of different ovarian stromal cell types. (G) The cell marker Lum’s location by 10× Visium sections and in situ hybridization. (H) The location of the cell marker Cyp17a1 in 10× Visium sections and immunofluorescence. (I) The cell marker Enpep’s location by 10× Visium sections and in situ hybridization. (J) The cell marker Aldh1a2’s location by 10× Visium sections and Immunohistochemistry.

### Fate transition of stromal cells along pseudotime trajectories and clear stromal cell classification

To dissect the fate determination of stromal cells throughout the investigated period, they were ordered along pseudotime trajectories according to gene expression (Figure 5A). We can see that the majority of cells on day 3, day 5, and day 7 were in the root state, and stromal cells began to be distinguished on day 7. This finding revealed that day 7 is the time point at which stromal cells differentiated. Cell pseudotime trajectories revealed 5 stromal cell states and 2 branch points (Figure 5B). Clusters 6 and 10 were in starting point of the root state (Figure 5C). When integrated analysis with the results of CytoTRACE and the specific cell markers were performed, it was found that cluster 6 may be the progenitor of stromal cells. As shown in Figure 5D, the cluster was characterized by the expression of genes associated with biological processes, such as “PI3K-Akt signaling pathway,” “focal adhesion,” and “protein processing in endoplasmic reticulum,” which were more highly expressed in state 1 cells, suggesting a naïve typical function of stromal cells. Genes encoding members of “lipid and atherosclerosis,” “MAPK signaling pathway,” and “FoxO signaling pathway” were among state 3, confirming the existence of steroid hormone production in stromal cells. State 5 was composed of genes encoding proteins involved in “Huntington disease,” “amyotrophic lateral sclerosis” and “oxidative phosphorylation,” which suggested the high functional state of these stromal cells. The genes Ptn and Gstm6 were mainly expressed in state 1, Enpep was mainly expressed in state 5, and Star was mainly expressed in state 3 (Figure 5E). Moreover, 4 gene sets with distinct patterns were identified (Figure 5F, Table S4), and the specifically represented genes of the 4 gene sets, namely, Sfrp1, Fhl2, Tmem100, and Cebpb, are shown in Figure 5G.

**Figure 5.**
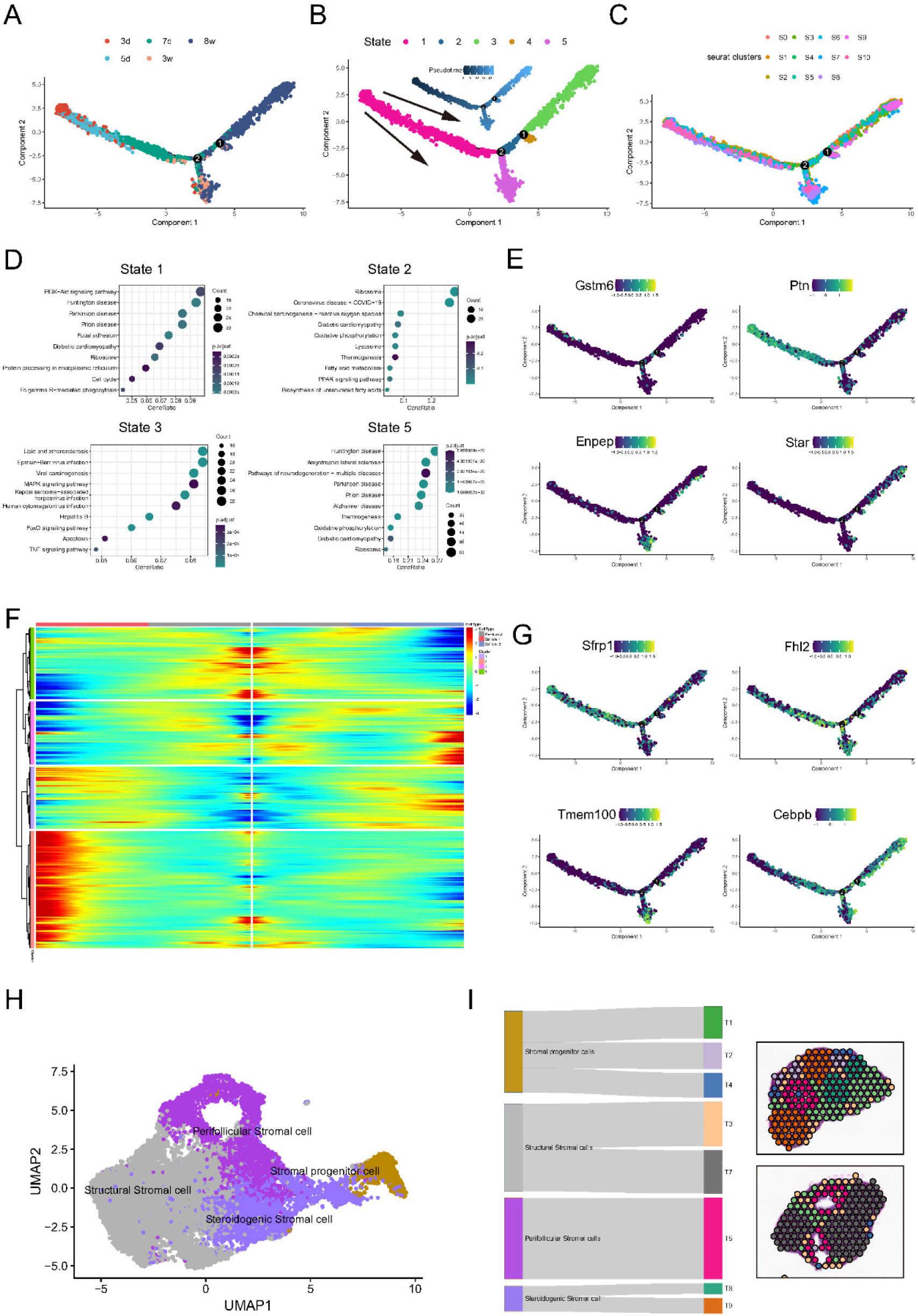
Identification of ovarian stromal cell subpopulations (A) Single-cell trajectories of ovarian stromal cells as a function of the developmental timeline. (B) Single-cell trajectories of the 3 stromal cell states through pseudotime analysis. (C) Single-cell trajectories of germ cells along the Seurat cluster. (D) GO analysis of states 1, 2, 3, and 5 through pseudotime. (E) Expression of representative genes in each set along single-cell pseudotime trajectories. (F) Heatmap representing the expression dynamics of 5 gene sets with increased expression or reduced expression at the follicle stage. (G) Expression profiles of the differentially expressed genes in the BEAM branch. (H) Four specific ovarian stromal cell populations. (I) MIA of the specific ovarian stromal cell populations mapping to ovarian tissue.

Combining all of the above results, we divided ovarian stromal cells into 4 subpopulations: structural stromal cells, perifollicular stromal cells, stromal progenitor cells, and steroidogenic stromal cells (Figure 5H). Multimodal intersection analysis (MIA) also confirmed the classification of stromal cells in 10× Visium sections (Figure 5I).

To further verify our ovarian stromal cell classification, we performed analysis with human ovary tissue around the follicle and mouse ovary at eleven time points between E11.5 and 8 weeks. Human ovarian tissues with follicles with diameters of 2 mm and 5 mm were included in the analysis (Figure 6A). The ovarian cells can be classified into five cell subpopulations according to the known cell markers, and the stromal cells can be classified into six clusters (Figure 6B). As shown in Figure 6C-E, ENPEP+ perifollicular stromal cells and CYP17A1+ steroidogenic stromal cells surrounded the human follicle. We then integrated our data with the ovaries at six time points between E11.5 and 1 day, and 11 cell types of ovarian cells were identified (Figure 6F-H). Among these cell types, stromal cells and GCs also accounted for the vast majority of the population (Figure 6I). The stromal cells were categorized into 10 clusters, and the majority of Aldh1a2+ stromal progenitor cells were noted at E11.5 and at the root of the pseudotime trajectories (Figure 6K-N).

**Figure 6.**
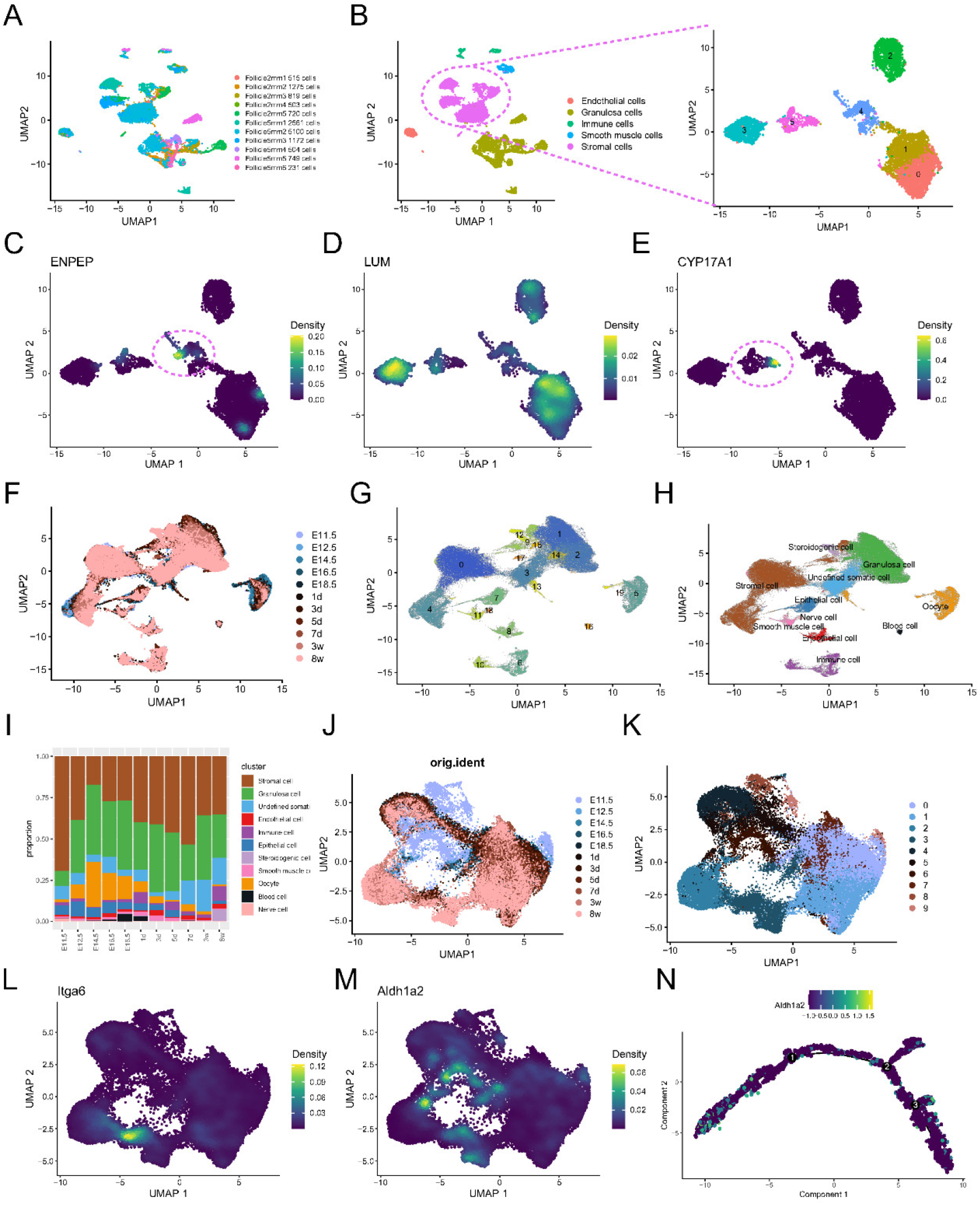
Identification of the ovarian stromal cell subpopulations in human ovary and early gonad of mouse (A) UMAP plot of ovarian cell colored by different human follicle samples; (B) UMAP plot of the classified 5 cell types and the ovarian stromal cell colored by cluster; (C) Feature plots of specific marker gene of Perifollicular Stromal cell, ENPEP in human follicle stroma; (D) Feature plots of specific marker gene of Structural Stromal cell, LUM in human follicle stroma; (E) Feature plots of specific marker gene of Steroidogenic Stromal cell, CYP17A1 in human follicle stroma; (F) UMAP plot of ovarian cells colored by the selected developmental time points (that is Stage); (G) UMAP plot of ovarian cells colored by cluster; (H) UMAP plot of 11 ovarian cell types. (I) Percentages of the 11 ovarian stromal cell clusters at E11.5, E12.5, E14.5, E16.5, E18.5, day 1, day 3, day 5, day 7, week 3 and week 8. (J) UMAP plot of ovarian stromal cell colored by the selected developmental time points (that is Stage) (K) UMAP plot of ovarian stromal cell colored by the clusters. (L and M) Feature plots of specific marker gene of Stromal progenitor cell, Itga6 and Aldh1a2; (N) Expression of Aldh1a2 along with pseudotime trajectories

Combining mouse and human single-cell data, the results confirmed our classification of ovarian stromal cells.

### Specific regulon networks of ovarian stromal cells during follicle development

To gain a deeper understanding of the functions of the above four groups of cells, we further investigated the regulon (transcription factor (TF)) activity of stromal cell-specific TFs using SCENIC. We identified several important TFs during follicle development. Based on 51 regulon activities with 5,481 filtered genes with default filter parameters, regulon activity was matched with developmental stages, and representative regulons are listed (Figure 7A). As shown, a series of TFs displayed a more dynamic pattern. For example, Maf, Wt1, Gata6, and Erg1 were active mainly at the early development stage (day 3 and day 5, Figure 7A-C). As follicle development continued, Foxo1, Runx1, Stat1, and Smarca4 seemed to be mostly activated at the 3-week and 8-week stages, when the follicle matured (Figure 7A and 7G). Since stromal cells began differentiating into different fates in cell pseudotime trajectories on day 7, we focused on this time point. Yy1, Jdp2, Cebpb, and Maff were active mainly on day 7 (Figure 7A and Figure 7D-E). The regulon activity of Bclaf1 was specifically turned off on day 7 (Figure 7F). These regulon activity patterns may provide new insight into ovarian stromal cell fate determination.

**Figure 7.**
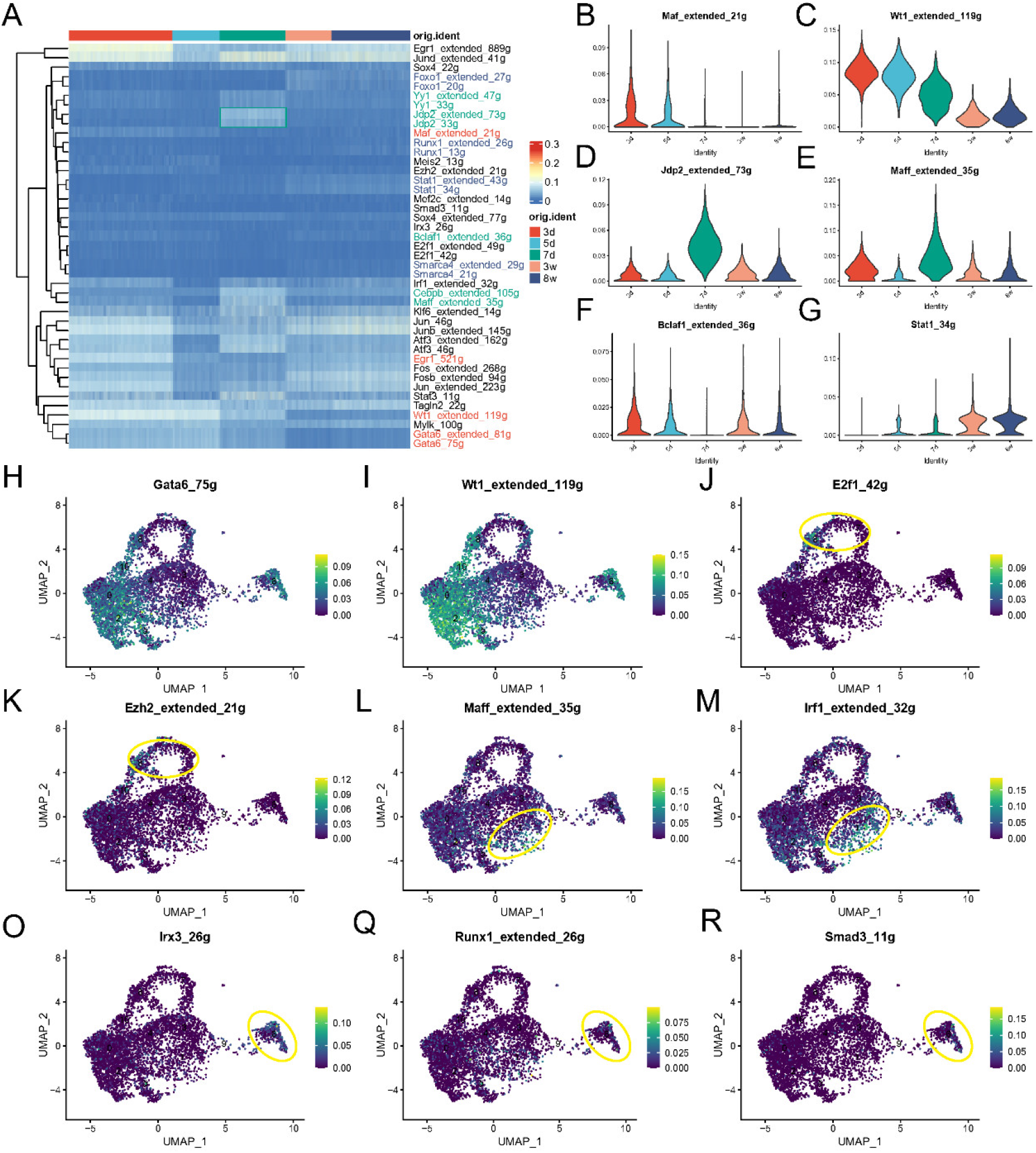
Specific regulon networks of ovarian stromal cell during follicle development (A) Heatmap of regulon activity analyzed by SCENIC with default thresholds. (B-G) Average regulon activity of Maf, Wt1, Jdp2, Maff, Bclaf1 and Stat1 that were stage specific in ovarian stromal cell during follicle development. (H-R) UMAP projection of average regulon activity of Gata6, Wt1, E2f1, Ezh2, Maff, Irf1, Irx3, Runx1, and Smad3 that were stage specific in ovarian stromal cell subpopulations.

Moreover, when we linked regulon activity with ovarian stromal cell subpopulations, we found that some TFs were significantly represented with specific ovarian stromal cell subpopulations. As shown in Figure 7H and I, Gata6 and Wt1 were active mainly in stromal cell subpopulations without perifollicular stromal cells. E2f1 and Ezh2 were active mainly in the perifollicular stromal cells (Figure 7J and K). TFs such as Maff and Irf1 were active mainly in steroidogenic stromal cells (Figure 7L and M). Irx3, Runx1, and Smad3 were active mainly in the stromal progenitor cell subpopulation (Figure 7O-R).

The generation of specific regulon networks of ovarian stromal cells identified several important TFs associated with different ovarian stromal cell subpopulations, helping us explore ovarian stromal cellular fate in vitro.

### Stromal-derived MK coculture promotes follicle development

As MK signaling is expressed throughout the total follicle development stage, we added midkine to coculture with follicles encapsulated in alginate to further confirm the role of outgoing ovarian stromal cell signaling in follicle development. As shown in Figure 8A, ovarian stromal cells were the major source of outgoing MK signaling, and GCs were the major target. Receptor–ligand analysis showed that Ncl, Sdc4, (Itga6+ Itgb1) and Lrp1 were major contributors to MK signaling (Figure 8B). We performed Mdk ISH analysis of mouse ovaries from day 3, day 5, day 7, week 3 and week 8. Mdk was mainly located in the ovary stroma and had the highest expression in day 5 mouse ovaries. As follicles develop, Mdk expression mainly occurs in the stroma around the growing follicle but decreases around the antral follicle or corpus luteum, suggesting that it may play a vital role in follicle development and maturation (Figure 8C). We then investigated the expression and localization of the ligands of Mdk. Both ISH and IHC showed that Ncl and Lrp1 were mainly expressed in GCs in growing follicles (Figure 8D and E). Next, we added purified MK to the culture medium to coculture with the early secondary follicle in vitro (Figure 8F). The results showed that midkine significantly promoted the increase in follicle growth (Figure 8G). These results confirmed that the role of ovarian stromal cell crosstalk with follicles promotes follicle development.

**Figure 8.**
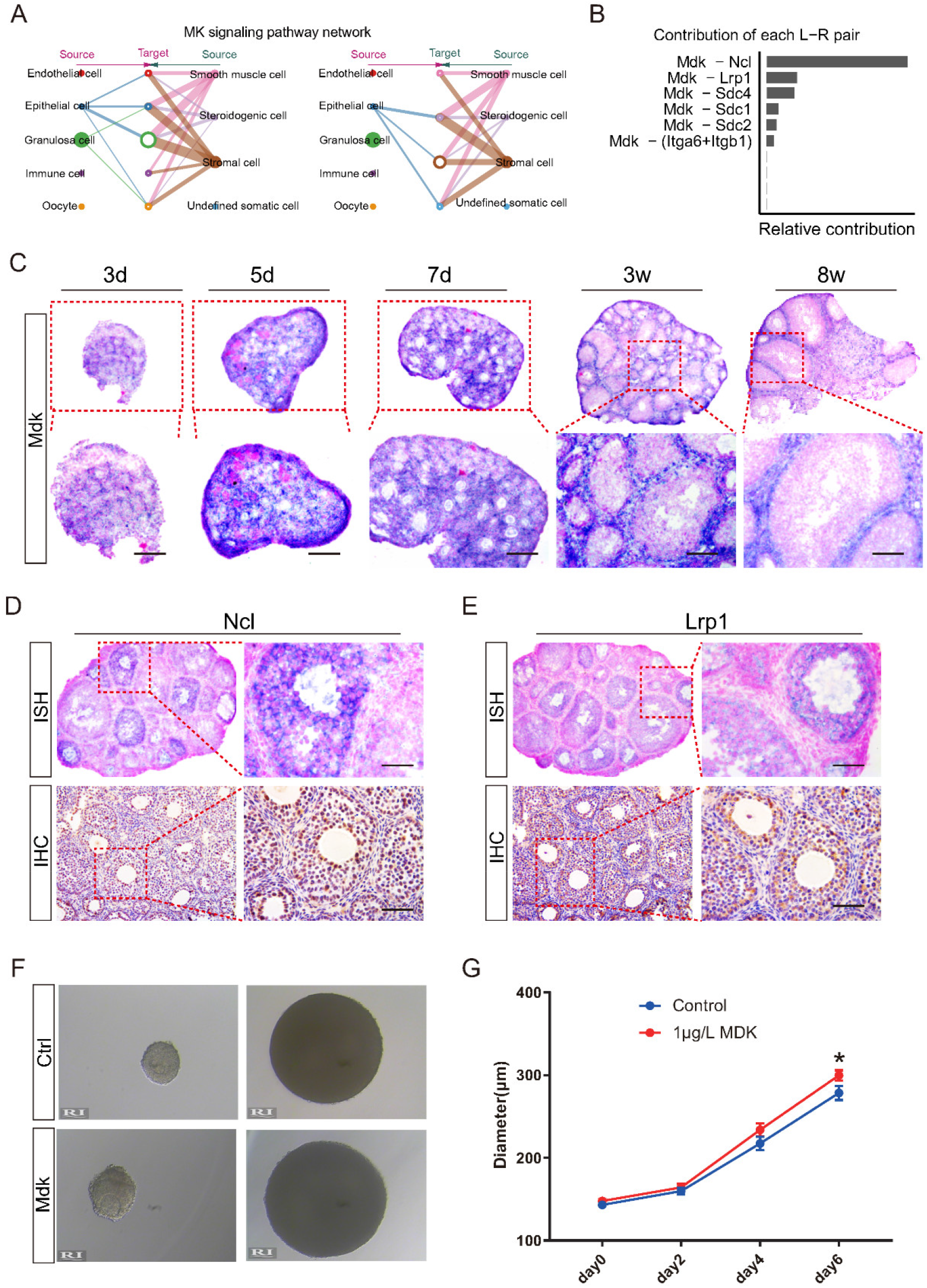
Stromal-derived Midkine co-culture promote follicle development (A) MK signaling pathway network among the nine ovarian cell types at week 3. (B) The contribution of each L-R pair of MK signaling pathway. (C) The represented picture of Mdk ISH in ovary section of day 3, day 5, day 7, week 3, and week 8 mice. Scale bar = 50μm. (D) The represented picture of Ncl of ISH and IHC analysis in ovary section of 3-week mice. Scale bar = 50μm. (E) The represented picture of Lrp1 of ISH and IHC analysis in ovary section of 3-week mice. Scale bar = 50μm. (F) The represented picture of follicle with/without addition of 500ng/l Mdk. (G) The diameter of follicles analysis in 6-day culture (*p < 0.05, Student’s t test).

### Perifollicular stromal cells can differentiate into theca-like cells in vitro

To study the role of stromal cell subpopulations in follicle development, we obtained perifollicular stromal cells by flow cytometry sorting using an ENPEP-PE antibody (Figure 9A). The ENPEP+ perifollicular stromal cells were more fibroblast-like than the ENPEP-cells (Figure 9B). Immunofluorescence analysis showed that ENPEP+ perifollicular stromal cells expressed ENPEP and COL1A1 but not FOXL2. The ENPEP-cells did not express ENPEP or COL1A1 and had high levels of FOXL2 expression (Figure 9C). qPCR analysis revealed that ENPEP+ perifollicular stromal cells had higher levels of Pdgfa, Enpep, and Mdk and lower levels of Amhr2 and Foxl2 (Figure 9D). During cell culture, ENPEP+ perifollicular stromal cells cannot proliferate when passaged to the third generation (P3), suggesting that ENPEP+ perifollicular stromal cells may differentiate in vitro. Immunofluorescence and qPCR analysis of the theca cell markers Star and Cyp17a1 confirmed that ENPEP+ perifollicular stromal cells differentiated into theca-like cells in long-term in vitro culture (Figure 9E-F). The results revealed the importance of ovarian stromal cell type classification. These results suggest that perifollicular stromal cells not only have a paracrine role but also have a differentiation function.

**Figure 9.**
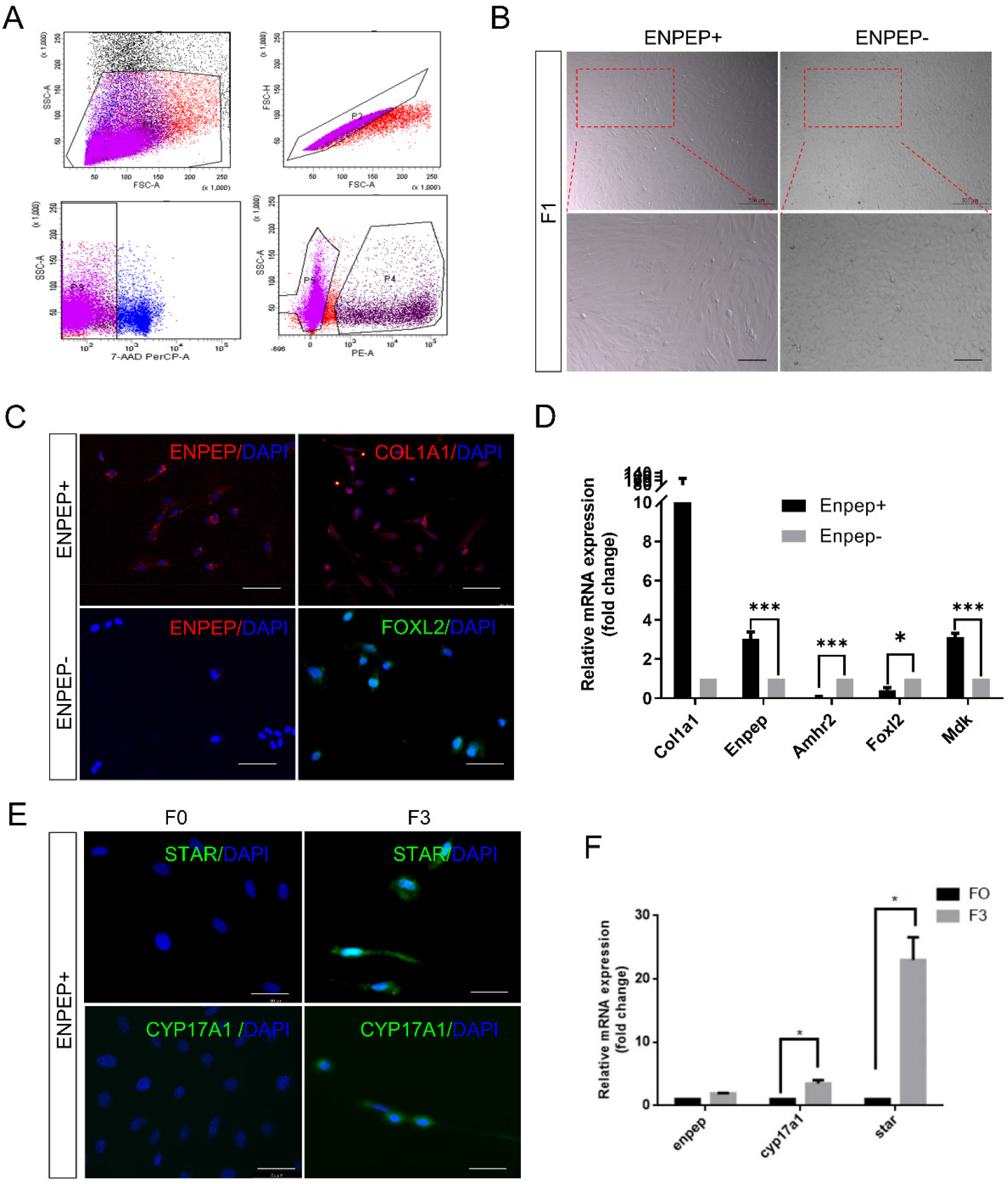
Perifollicular stromal cell can differentiate into theca-like cell (A) Perifollicular stromal cell sorting strategy using ENPEP-PE antibody. (B) The represented pictures of ENPEP+ and ENPEP-cells after sorting in P1 generation. Scale bar = 500μm, 200μm. (C) Immunofluorescence of ENPEP, COL1A1, and FOXL2 confirmed the characteristic of perifollicular stromal cell. Scale bar = 50μm. (D) qPCR analysis of Col1a1, Enpep, Amhr2, and Mdk in ENPEP+ and ENPEP-cells after sorting (*p < 0.05, ***p < 0.001, Student’s t test). (E) Immunofluorescence of STAR and CYP17A1 confirmed that perifollicular stromal cell differentiates into theca-like cell in vitro. Scale bar = 50μm. (F) qPCR analysis of Enpep, Cyp17a1, and Star of P0 and P3 perifollicular stromal cell (*p < 0.05, Student’s t test).

## Discussion

Ovary and follicle development is a complex process in mammals, and a full understanding of its regulation requires the integration of multiple types and spatial location of data collected from various cell types in the ovary. Here, we performed scRNA-seq and spatial transcriptomics analysis of all the cell types within the mouse ovaries at five different developmental stages (from the primordial follicle activation stage to the mature follicle stage, day 3 to 8 weeks) to provide new and comprehensive insights into the regulation of ovary and follicle development in mice. In this study, the niche cells of follicles, such as stromal cells, GCs, epithelial cells, endothelial cells, and immune cells, were identified. Furthermore, our work revealed unique characteristics in the transcriptional profiles and cell communication between stromal cells and other cell types in each developmental stage. Moreover, we identified four clusters of stromal cells that were incompletely characterized and categorized previously: structural stromal cells, perifollicular stromal cells, stromal progenitor cells, and steroidogenic stromal cells. Structural stromal cells are distributed throughout the ovary and mainly perform supportive functions. Perifollicular stromal cells are distributed around growing follicles and not only regulate follicular development through secretory signals but can also differentiate and replenish theca cells. Stromal progenitor cells are mainly located in the ovarian cortex and are the progenitor of all stromal cell types. Steroidogenic stromal cells were distributed in the ovarian medulla and synthesize steroid hormones. These identified cell types and the stage-specific gene expression in ovary development offer valuable information for future functional studies.

With developments in scRNA and spatial transcriptomics technologies to complement detailed imaging studies, we have the ability to better characterize the ovarian cell populations and to refer to them with more precise names as we understand their individual roles in physiological and pathological processes. Most of the previous studies on follicle development focused on the follicle body and ignored the function of stromal cells. Our study found that the stroma is crucial for follicular development, and it is far from enough to focus on the follicle alone. The subpopulation of ovarian stromal cells is complex, and regional differences likely influence the distribution and subtypes of stromal cells. The multiple populations of ovarian stromal cells are incompletely characterized, but we have identified four types of ovarian stromal cells with the combination of scRNA with spatial transcriptomics technology. Single-cell RNA-sequencing experiments have already made progress in identifying major ovarian cell types, transition stages, and markers for cell identification. However, data about ovarian stromal cells remain elusive and comparatively scarce in previous studies. Recent scRNA data from mice and humans have shown heterogeneity of ovarian stromal cells, but these studies mainly focused on GCs and oocytes^16–21^. Single-cell sequencing fails to provide spatial information of cell populations, which leads to incomplete characterization of ovarian stromal cells.

Follicles maintain contact with the supporting stromal cells, which provide the local biochemical control pathways that trigger the activation of follicle growth. Stroma-to-follicular paracrine signaling is very important for follicle development. To explore cell-to-cell communication between ovarian stromal cells and other cell populations of the ovary, it is critical to elucidate the impact of stromal cells on the perifollicular microenvironment. In this study, we mapped the secretory signaling networks between various cell types within the ovary and revealed the role of ovarian stromal cells as the major outgoing paracrine signaling. The secretory signals that we identified such as MDK provide new insight into the role of stromal cells in the follicle microenvironment. Hence, the impact of coculture could be significantly enhanced by integrating stromal cells into current three-dimensional culture systems. Previous studies have also shown that the bone morphogenetic proteins BMP-4 and BMP-7, which are expressed by stromal cells, have been linked to the primordial-to-primary follicle transition^22, 23^. The paracrine signaling involved in these processes is hypothesized to be a complex time-dependent synergy of unidentified factors.

In this study, we found a subpopulation of ovarian stromal cells surrounded by growing follicles and adjacent to theca cells, named ovarian perifollicular stromal cells. Ovarian perifollicular stromal cells not only send secretory signals (such as MDK) but can also differentiate into steroidogenic stromal cells, which express the theca cell markers Star and Cyp17a1. Our findings support that theca cells are recruited from surrounding stromal cells and identify a special subpopulation of stromal cells. Theca cells function in a diverse range of necessary roles during folliculogenesis; they synthesize androgens, provide crosstalk with GCs and oocytes during development, and provide structural support for the growing follicle as it progresses through the developmental stages to produce a mature and fertilizable oocyte^24^. Theca cells were previously thought to be recruited from surrounding stromal tissue by factors secreted from an activated primary follicle^4^. The precise origin and identity of these recruiting factors are currently not clear^24^. These specialized cells have long been thought to originate from fibroblast-like stromal cells. The putative undifferentiated progenitor theca cells do not express LH receptors (LHRs) or steroidogenic enzymes and are therefore not LH responsive, showing that initiation of theca cell differentiation is gonadotropin independent^24^. Recently, studies have also shown that autologous transplantation of thecal stem cells restores ovarian function in nonhuman primates^25^. The thecal stem cells used were total ovarian stromal cells, not a single-cell population. Our study identified and obtained functional perifollicular stromal cells by cell sorting that can differentiate into theca cells. This finding sheds new light on the development of stem cell therapy for premature ovarian insufficiency (POI).

Stromal cells are assumed to arise from a population of unspecialized mesenchymal stem cells, but their origins and physiological roles remain elusive, and more compelling data are needed. We have identified that Aldh1a2 is a marker gene of ovarian stromal progenitor cells and is expressed mainly in ovarian surface epithelial cells and stromal cells in the ovarian cortex. Aldh1a2+ ovarian stromal progenitor cells were mainly at the origin of stromal cell development in the pseudotime trajectories. We also noticed that the Aldh1a2+ ovarian stromal progenitor cell population decreased over time, which suggests that the attenuation of ovarian function is not only a decline in the follicle pool but also a decline in ovarian stromal progenitor cells. Aldehyde dehydrogenase 1 (ALDH1) is a marker of normal tissue stem cells, where it is involved in self-renewal, differentiation, and self-protection. Studies have demonstrated Aldh1a1- and Aldh1a2-expressing stem cell niches in ovarian surface epithelial cells and subsurface regions^26, 27^.

Ovarian stromal abnormalities are an important cause of many clinical ovarian dysfunction diseases, such as polycystic ovary syndrome (PCOS). PCOS is a common endocrine disorder in women and is characterized by oligo- or anovulation, hyperandrogenism, and ovarian polycystosis. Cortical thickening, stromal hyperplasia, and cystic antral follicles were found in the ovaries of PCOS patients, accompanied by significant increases in ovarian volume, stromal volume, and increased stromal blood flow velocity^28^. Superovulation therapy rescues abnormal stroma morphology, resulting in the acquisition of mature oocytes in some PCOS patients. Abnormalities in stromal cells are an important cause of PCOS, but the mechanism is unclear. Angiopoietin-like protein (Angptl-4)^29^ and TWEAK^11^ are thought to be involved in the pathogenesis of PCOS and are two major signaling pathways through which ovarian stromal cells communicate with other cell types. Moreover, our study showed that ovarian perifollicular stromal cells grow along with follicles and can differentiate into theca-like cells; thus, the cause of abnormal stroma in PCOS may be the dysfunction of ovarian perifollicular stromal cells. The abnormal proliferation of ovarian perifollicular stromal cells can lead to stromal hyperplasia, and increased differentiation to theca-like steroidogenic stromal cells can result in hyperandrogenism. Therefore, our classification of ovarian perifollicular stromal cells provides new insight into the occurrence and treatment of PCOS.

In summary, this work provides new insights into the crucial features of mouse follicle development, especially stromal cells. Our study paves the way for understanding the molecular regulation of ovarian stromal cells in follicle development. It also lays a solid foundation for ovarian stromal cell classification and origin identity. We obtained functional perifollicular stromal cells and identified the vital secreted factors that promote follicle development in vivo and in vitro. More importantly, we have uncovered the reciprocal relationship between the signaling pathways of follicles and their niche cells, which will provide a valuable resource for further optimizing and improving the efficiency of follicle culture in vitro. Our understanding of stromal cell subpopulations will help us better understand stromal cell hormone production and responsiveness, pathological ovarian stromal changes, such as polycystic ovary syndrome or premature ovarian insufficiency, and stromal contributions to artificial ovary technology.

## Materials and methods

### Animals and preparation of cell suspensions

C57BL/6J mice were purchased for scRNA-seq and ovarian section staining from the Animal Experiment Center of Nanjing Medical University (Nanjing, China). Ovaries were dissected from female mice at 3 days, 7 days, 3 weeks, and 8 weeks, and we also included the 5-day data from published reports^30^ and 10-month ovary data (data not shown). The selected 5 time points for sequencing represent the key stage of a series of cellular events involving ovary and follicle development. Moreover, 7 days, 3 weeks, and 8 weeks ovaries were collected and embedded with OCT. Ten-micron sections were taken for the 10× Visium experiment.

To obtain single-cell populations, isolated ovaries were cut into small pieces and incubated in 0.25% trypsin (Gibco, Grand Island, NY, USA) and collagenase II (0.2%, Sigma–Aldrich, C2-22-1G, Darmstadt, Germany) for 6 to 8 min at 37°C. Tissues were disaggregated with a pipette to generate single cells, and the solution was filtered through 40-μm cell strainers (BD Falcon, 352340, USA) and washed 2 times with PBS containing 0.04% BSA. Cell viability was acceptable after staining in 0.4% Trypan Blue; it was above 80%, and the cell concentration (1,000 cells/μL) was checked to meet the requirements for sequencing, as was the single-cell rate (no connected cells observed during cell counting).

### Ethics statement

The mice were maintained under specific pathogen-free conditions in a controlled environment at a temperature of 20±2°C and a humidity of 50–70% on a 12:12-h light:dark cycle, with food and water provided ad libitum. Animal care and experimental procedures were performed in accordance with the guidelines of the Experimental Animal Management Committee (Jiangsu, China) and were approved by the Ethics Review Board for Animal Experiments of the Affiliated Drum Tower Hospital of Nanjing University Medical School. All applicable institutional and/or national guidelines for the care and use of animals were followed.

### Single-cell libraries and sequencing

Single-cell suspensions of ovary cells were captured on a 10× Chromium system (10× Genomics). Then, single-cell mRNA libraries were generated using single-cell 3’ reagent V3 kits according to the manufacturer’s protocol. After the generation of GEMs, reverse transcription reactions were barcoded using a unique molecular identifier (UMI), and cDNA libraries were then amplified by PCR with appropriate cycles. Subsequently, the amplified cDNA libraries were fragmented and then sequenced on an Illumina NovaSeq 6000 (Illumina, San Diego). Furthermore, the “mkfastq” module of Cell Ranger was used to produce FASTQ files with raw base call (BCL) files as input generated by Illumina sequencer alignment.

### 10× Visium processing and sequencing

The ovary tissues obtained were cut at 10-μm thickness, mounted onto corresponding Capture Areas on the Visium Spatial Tissue Optimization Slide. Next, the tissue was dehydrated with isopropanol for 1 min followed by staining with hematoxylin and eosin. Slides were mounted in 80% glycerol and brightfield images were taken on 3D HISTECH Pannoramic MIDI FL, whole-slide scanner at 40× resolution. After permeabilized, and cellular mRNA is captured by the primers on the gene expression spots. All the cDNA generated from mRNA captured by primers on a specific spot share a common Spatial Barcode. Libraries are generated from the cDNA and sequenced and the Spatial Barcodes are used to associate the reads back to the tissue section images for spatial gene expression mapping. Visium Spatial Gene Expression libraries comprise standard Illumina paired-end constructs that are flanked with P5/P7, necessary for binding to the Illumina flow cell.

### scRNA-seq data processing and analysis

The Cell Ranger “count” pipeline (version 3.1.0) was applied with the FASTQ data produced to map the mouse reference genome (version mm10) and process feature–barcode matrixes with default parameters. To prevent potential bias during subsequent bioinformatics analysis, the number of captured cells in each sample was set to 6,000 with the parameter “force-cells.” The data matrixes were then loaded in R (version 4.1) using the Seurat package (version 4.1.0)^31^. The Seurat object was created based on 2 filtering parameters of “min.cells = 5” and “low.thresholds = 200,” and the exorbitant number of unique genes detected in each cell (i.e., “nFeature_RNA”) was adjusted in each sample to eliminate the empty drops and dying cells and potential doublets/multiplets from subsequent analyses. Then, the multiple samples were processed with “harmony”^32^. Following the normalization and scaling steps, using the UMAP (a visualized method for cell clustering of high-dimensional transcriptomic data) technique^33^, a series of commands were executed to visualize cell clusters, including the “RunUMAP” function with a proper combination of the “resolution” and “dims.use,” “FindNeighbors” and “FindClusters” functions to conduct cell clustering. To identify canonical cell cluster marker genes, the “FindAllMarkers” function was used to identify conserved marker genes in clusters with default parameters.

### 10×Visium data preprocessing

Raw FASTQ files and histology images were processed by sample with the SpaceRanger software version 1.2.0, which uses STAR for genome alignment, against the mm10 reference genome. We processed the unique molecular identifier (UMI) count matrix using the R package Seurat and normalized the data by SCTransform^34^ to account for variance in sequencing depth across data points. Top variable genes across single cells were identified using the method described in Macosko et al^35^. Cells were clustered based on a graph-based clustering approach and were visualized in 2-dimension using UMAP.

We use SPOTlight^36^ to integrate 10× Visium with scRNA-seq data to infer the location of cell types and states within a complex tissue. Further, Multimodal intersection analysis (MIA) was according to the method described in Moncada et al^37^.

### Pseudotime analysis of single-cell transcriptomes

Cell lineage trajectories were constructed according to the procedure recommended in the documentation for Monocle2^38^ (http://cole-trapnell-lab.github.io/monocle-release/docs), a novel unsupervised algorithm, which reordered the cells to maximize the transcriptional similarity based on their differentiation progress, and at the same time, distinguished genes that are activated early and later in differentiation, which contribute to dissecting cell differentiation fate, also termed “pseudotime analysis.” With the gene count matrix as input, the new dataset for Monocle object was created, and the functions “reduceDimension” and “orderCells” were carried out to generate the cell trajectory based on pseudotime. In particular, the ordering genes were differentially expressed genes between clusters in each cell type calculated by the “differentialGeneTest” function in Monocle. In addition, the “BEAM” function was used to calculate the differentially expressed genes at a branch point in the trajectory, and genes with “qval < 1 × 10e-4” are shown with a heatmap. Moreover, the root state (that is, a prebranch in the heatmap) was set and adjusted following consideration of the biological meanings of different cell branches.

### GO and KEGG enrichment analyses of the gene set

For a large gene set with more than 100 genes, the GO and KEGG enrichment analyses were performed with ClusterProfiler^39^, an R package in Bioconductor, to detect the gene-related biological process, and signaling pathways with a threshold value of “pvalueCut off = 0.05” and top terms were displayed.

### Regulon activity of transcription factors with SCENIC

The SCENIC algorithm was developed to assess the regulatory network analysis of TFs and to discover regulons (that is, TFs and their target genes) in individual cells^40^. Following the standard pipeline, the gene expression matrix with gene names in rows and cells as columns was input into SCENIC (version 0.9.1). The genes were filtered with default parameters, and finally, 9,208 genes were available in the RcisTarget database, the mouse-specific database (mm10) that is used as the default in SCENIC. The coexpressed genes for each TF were constructed with GENIE3 software, followed by Spearman’s correlation between the TF and the potential targets, and then the “runSCENIC” procedure assisted in generating the GRNs (also termed regulons). Finally, regulon activity was analyzed by AUCell (Area Under the Curve) software.

### Digoxigenin (DIG)-labeled RNA probe in situ hybridization

The ovary sections were baked at 37°C and fixed in 4% PFA at room temperature for 20 min. After acetylation, the preheated probes (Table S5) were evenly added to the sections and hybridized at 60°C overnight. The sections were washed with post-hybridization washing solution, MABT solution and RNA washing solution and blocked at room temperature for 1 h. The AP solution of anti-DIG was diluted at 1:2500, and the antibody was incubated with the sections at 4°C overnight. We cleaned the sections with MABT solution at room temperature 4 times and with NTM once, added chromogenic solution (Biyotime Biotechnology, C3206) to the sections, and then stained the nuclei after cleaning. The sections were treated with double-distilled water, a series of gradient alcohol and xylene after dyeing, and finally were mounted with SlowFade® Gold antifade reagent (Life Technologies).

### Follicle culture

Two-layered secondary follicles were mechanically isolated from 12-day-old female F1 hybrids (C57BL/6J×DBA/2J) by using insulin-gauge needles in L15 media (Invitrogen, Carlsbad, CA) containing 1% fetal calf serum (FCS)^41^. Individual follicles were maintained in minimal essential medium (aMEM)/1% FCS at 37°C and 5% CO_2_ for 2 h before encapsulation. Only those follicles displaying the following characteristics during the 2-h preincubation period were selected for encapsulation and culture: 1) diameter of 100–130 μm, 2) intact nature with some attached fibroblast-like theca cells, and 3) a visible immature oocyte that was round and centrally located within the follicle. Selected follicles were then encapsulated into 0.25% alginate beads. The alginate beads were left in the encapsulation solution for 2 min to cross-link the alginate and were then rinsed in culture media (aMEM with 10 mIU/ml recombinant FSH, 3 mg/ml BSA, 1 mg/ml bovine fetuin, 5 μg/ml insulin, 5 μg/ml transferrin, and 5 ng/ml selenium). Alginate beads containing a single follicle were plated at one follicle per well in 96-well plates in 100 μl of culture media. Encapsulated follicles were cultured at 37°C in 5% CO_2_ for 6 days.

### Immunohistochemistry

Ovarian samples were removed and fixed in 10% formalin solution at room temperature overnight; dehydrated in 70%, 80%, 90%, 95% and 100% ethanol solutions; and then embedded in paraffin^42^. Sample sections of 5 μm in thickness were deparaffinized and dehydrated with <u>xylene</u> and an ascending series of alcohol solutions. After antigen retrieval, the sections were incubated with the primary antibody (Table S5) at 4°C overnight and were washed with TBST solution at room temperature 3 times. Then, the sections were incubated with biotin-labeled secondary antibody at room temperature for 30 min, washed with TBST solution, and developed using a DAB peroxidase substrate kit (Zsbio, Beijing, China).

### Immunofluorescence

Sections of ovaries were deparaffinized and rehydrated with xylene and an alcohol gradient, and heat-induced antigen retrieval was performed in 10 mM sodium citrate buffer (pH 6.0)^43^. After permeabilization with 1% Triton X-100 in PBS, the sections were blocked with 3% BSA in PBS for 60 min at room temperature. The sections were incubated with primary antibodies (Table S5) diluted in blocking solution at 4°C overnight and washed with TBST solution 3 times for 5 min each time. We incubated the sections with secondary antibodies for 45 min and then counterstained them with DAPI (Life Technologies, Carlsbad, USA) for 10 min. The sections were mounted using SlowFade® Gold Antifade Reagent (Life Technologies) and examined with a confocal laser scanning microscope (Leica, Wetzlar, Germany).

### Flow cytometric analysis

The ovaries of 3-week-old mice were washed with 1× PBS 3 times, digested at 37°C with 0.1% trypsin/EDTA (Sigma, T8003) and 0.8 mg/ml type I collagenase (Worthington, LS004197), harvested as single-cell suspensions and resuspended in 1× PBS. The cells were incubated with phycoerythrin (PE)-conjugated Enpep antibody at 4°C for 60 min and then sorted by FACScan flow cytometry (Becton, Dickinson and company, Franklin Lakes, NJ).

### Statistical analysis

The data are presented as the mean ± SD and were obtained from 3 independent experiments. The statistical analysis was performed with GraphPad Prism software (version 9.0), and significant differences were determined with a Student’s unpaired t test. Significant and highly significant differences were set at P < 0.05 and P < 0.01, respectively.

## Data sharing statement

Please contact Dr. Sun and Dr. Yan to request all data and reagents described in this article.

## Acknowledgement

We thank all researchers whose work was relevant to but not cited in this article due to limited space.

---

**Figure S1.**
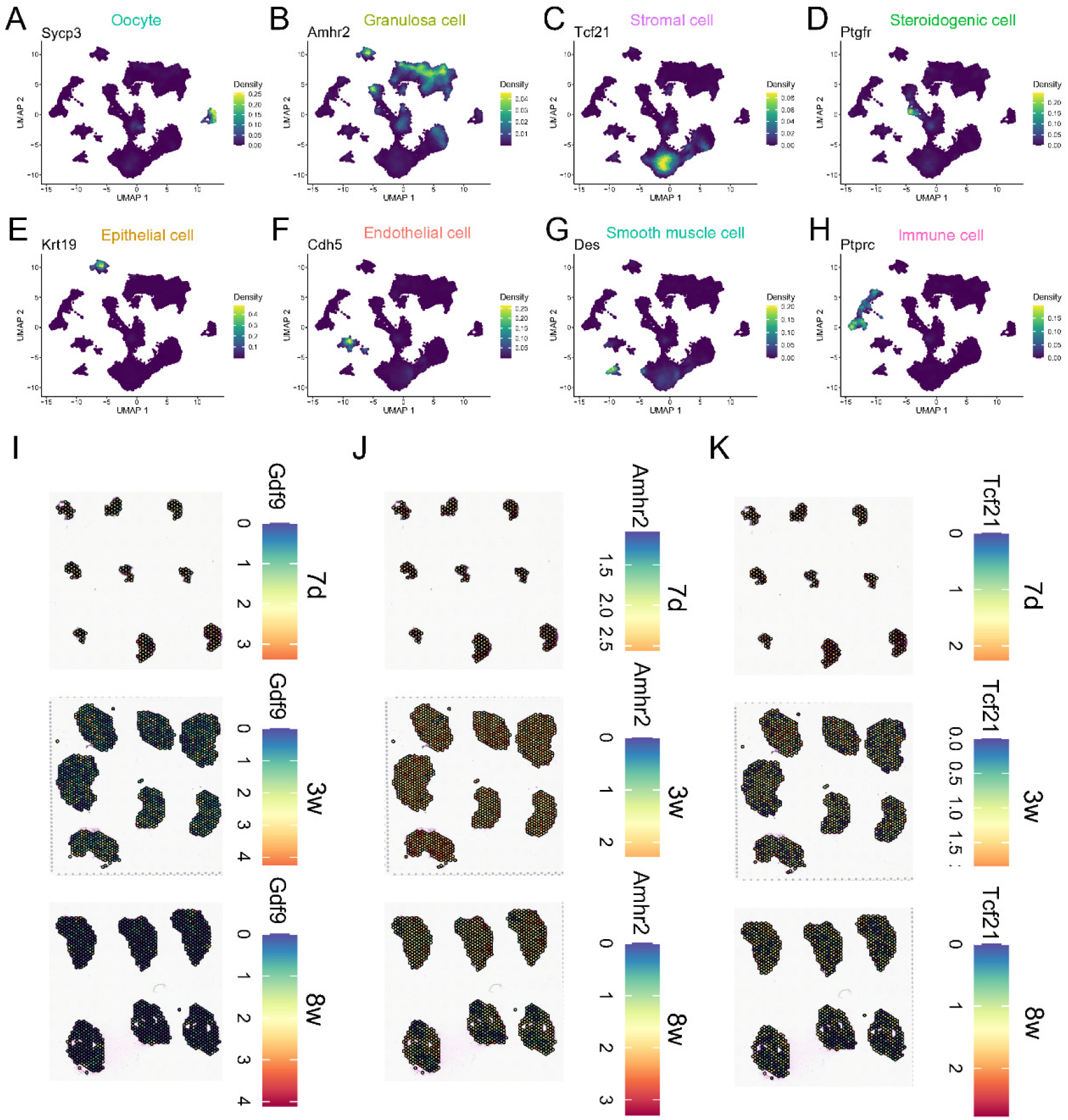
Specific marker genes of different ovarian cell types expression and location (A-H) Feature plots of specific marker genes of different ovarian cell types: Stromal cell, Granulosa cell, Oocyte, Endothelial cell, Epithelial cell, Immune cell, Smooth muscle cell, Steroidogenic cell. (I-J) The expression and location of specific marker genes of oocyte, granulosa cell, and stromal cell in 10× visium section.

**Figure S2.**
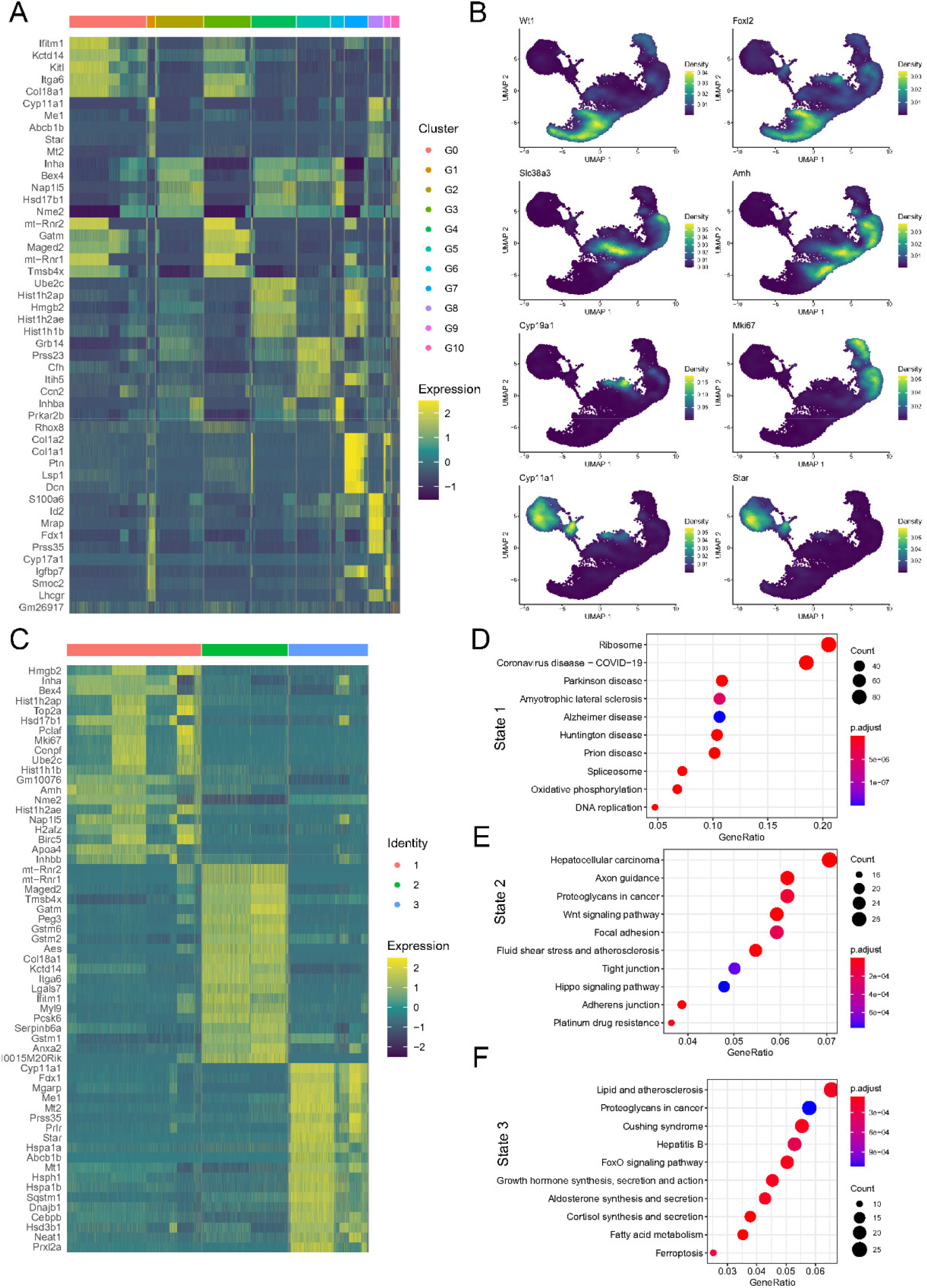
Transcriptional signature of granulosa cells (A) Heatmap of top 5 marker genes of the granulosa cell clusters. Top 10 marker genes in each cluster are shown in Table S2. (B) Feature plots of specific marker genes of different stromal cell types. (C) Heatmap of top 20 marker genes of single-cell trajectories of the 3 granulosa cell states. (D-E) Go analysis of top 100 genes of single-cell trajectories of the 3 granulosa cell states.

**Figure S3.**
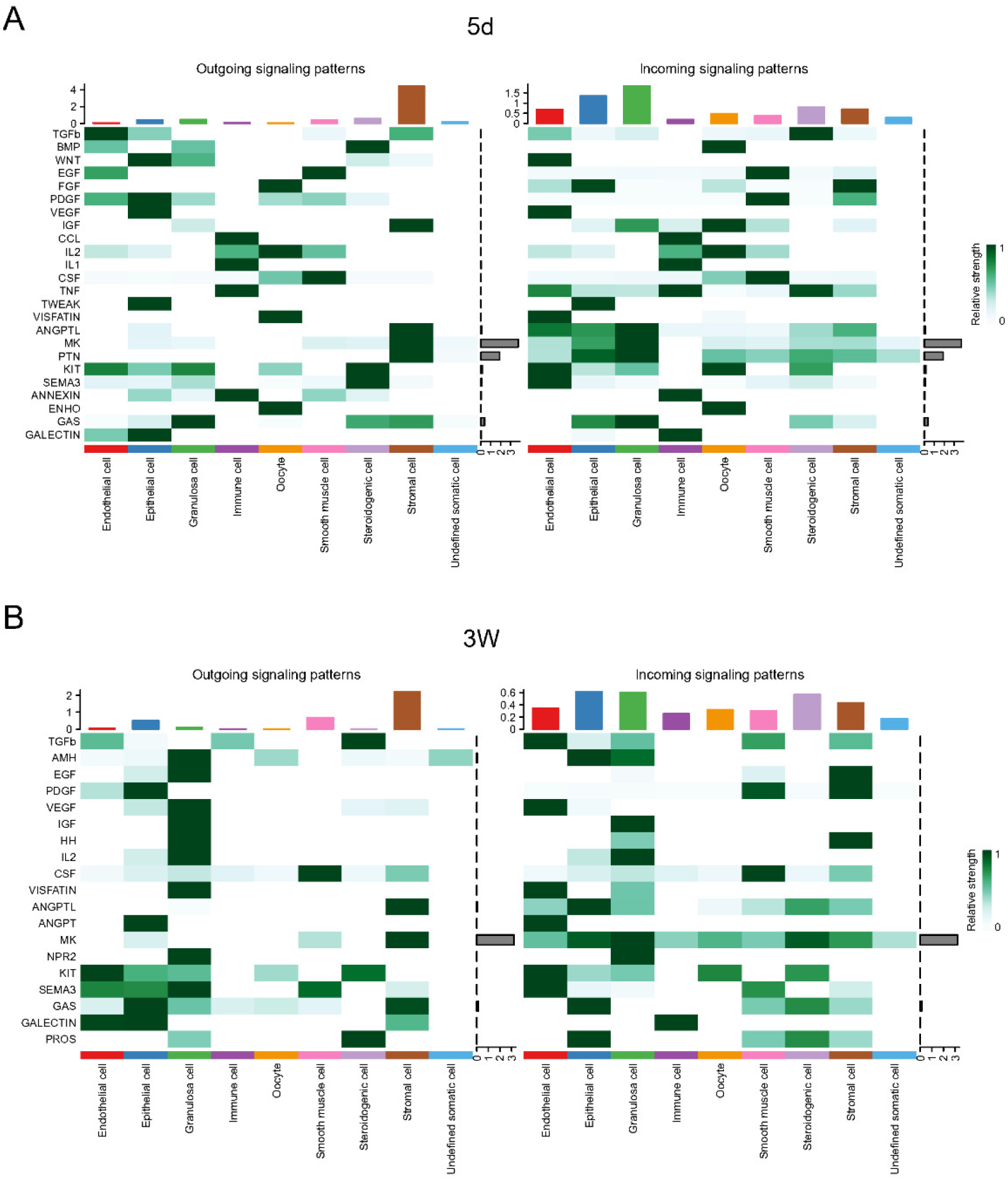
The outcoming and incoming signaling pattern of the nine ovarian cell types in the vital follicle development time pionts. (A) Outcoming and incoming signaling pattern of the nine ovarian cell types in the 5-day ovary mice. (B) Outcoming and incoming signaling pattern of the nine ovarian cell types in the 3-week ovary mice.

**Figure S4.**
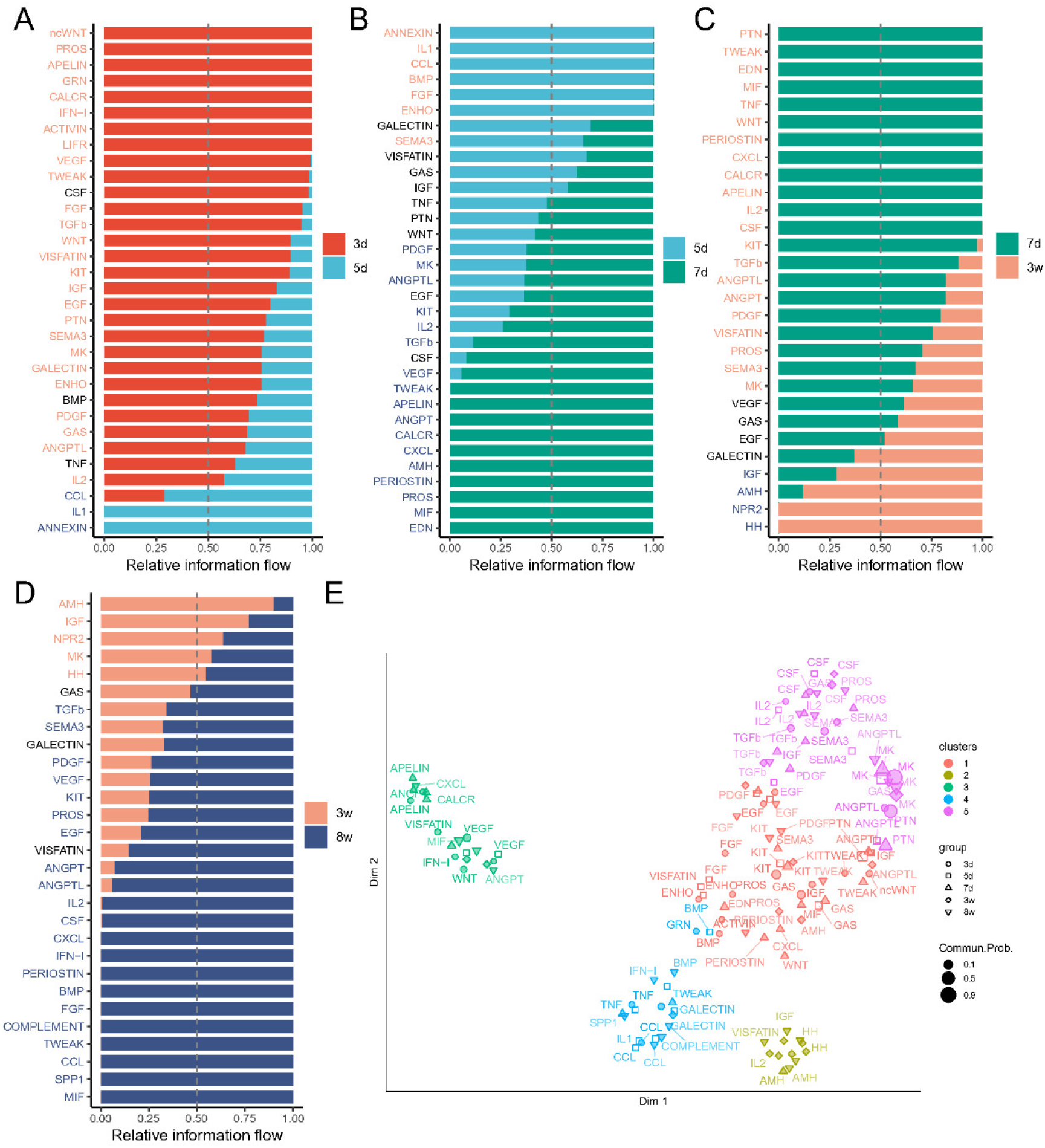
The relative information flow between the five time points of mice ovary. (A) The relative information flow between day 3 and day 5 mouse ovary. (B) The relative information flow between day 5 and day 7 mouse ovary. (C) The relative information flow between day 7 and 3 week mouse ovary. (D) The relative information flow between 3 week and 8 week mouse ovary. (E) Identify signaling networks of time points with larger (or less) difference based on their functional similarity

