## Supplementary material for "Temporal and spatial dynamics mapping reveals follicle development regulated by different stromal cell populations": Table S5

Supplemental Table 5 Information of antibody, primer sequences, and reagents

Antibody information

| Protein name | Manufacture (catalogue number) | Applications  (working dilution) |
| --- | --- | --- |
| Nucleolin | abcam(ab129200) | IHC(1:1000) |
| Lrp1 | abcam(ab92544) | IHC(1:1000) |
| Aidh1a2 | abcam (ab7574) | IHC(1:2000) |
| Cyp17a1 | Proteintech(14447-1-AP) | IHC(1:1000) |
| Enpep | Santa Cruz Biotechnology(sc-52444) | FCA(1 μg per 1x10^6 cells) |

Primer sequences

| Gene name | Sequences (5’-3’) | Application |
| --- | --- | --- |
| *MDK* | 5'>AATTAACCCTCACTAAAGGGATGCAGCACCGAGGCTTCTT<3' | ISH |
|  | 5'>TAATACGACTCACTATAGGGTTAGTCCTTTCCTTTTCCTTTCTTG<3' | ISH |
| *Enpep* | 5'>AATTAACCCTCACTAAAGGGATGAACTTTGCAGAGGAAGAGC<3' | ISH |
|  | 5'>TAATACGACTCACTATAGGGTCTGCAGCCTGGATCACCAC<3' | ISH |
| *Lum* | 5'>AATTAACCCTCACTAAAGGGATGAATGTATGTGCGTTCTCTCT<3' | ISH |
|  | 5'>TAATACGACTCACTATAGGGTTAGTTAACGGTGATTTCATTTGCTAC<3' | ISH |
| *Nucleolin* | 5'>AATTAACCCTCACTAAAGGGATGGTGAAGCTCGCAAAGG<3' | ISH |
|  | 5'>TAATACGACTCACTATAGGGCATCCTCTGAGGCAGGAGCA<3' | ISH |
| *Lrp1* | 5'>AATTAACCCTCACTAAAGGGACTATGGATGCCCCTAAAACTTG<3' | ISH |
|  | 5'>TAATACGACTCACTATAGGGATCTACTGGCTCATTCTTGGC<3' | ISH |
| *Foxl2* | 5'>ACAACACCGGAGAAACCAGAC<3' | q-PCR |
|  | 5'>CGTAGAACGGGAACTTGGCTA<3' |  |
| *Amhr2* | 5'>GGGGCTTTGGACACTGCTT<3' | q-PCR |
|  | 5'>GTCTCGGCATCCTTGCATCTC<3' |  |
| *Gapdh* | 5'>AGGTCGGTGTGAACGGATTTG<3' | q-PCR |
|  | 5'>TGTAGACCATGTAGTTGAGGTCA<3' |  |
| *Cyp17a1* | 5'>GCCCAAGTCAAAGACACCTAAT<3' | q-PCR |
|  | 5'>GTACCCAGGCGAAGAGAATAGA<3' |  |
| *StAR* | 5'>ATGTTCCTCGCTACGTTCAAG<3' | q-PCR |
|  | 5'>CCCAGTGCTCTCCAGTTGAG<3' |  |
| *Ptn* | 5'>ATGTCGTCCCAGCAATATCAGC<3' | q-PCR |
|  | 5'>CCAAGATGAAAATCAATGCCAGG<3' |  |
| *Mdk* | 5'>GAAGAAGGCGCGGTACAATG<3' | q-PCR |
|  | 5'>GAGGTGCAGGGCTTAGTCA<3' |  |
| *Col1a1* | 5'>GCTCCTCTTAGGGGCCACT<3' | q-PCR |
|  | 5'>GCTCCTCTTAGGGGCCACT<3' |  |
| *Enpep* | 5'>ATAGTGGGACTTTCTGTGGGT<3' | q-PCR |
|  | 5'>GGTCGTAGTGAACTGGATTGATG<3' |  |

Reagents

| Reagent | Source | Identifier |
| --- | --- | --- |
| DAB peroxidase substrate kit | Zsbio | ZLI-9018 |
| SlowFade® Gold Antifade Reagent | Beyotime | P0126-25ml |
| Chromogenic solution | Biyotime Biotechnology | C3206 |
| Trypsin/EDTA | Sigma | T8003 |
| Type I collagenase | Worthington | LS004197 |
| MEM Alpha(1X) | Gibco | 32561-037 |
| Recombinant Human Follitropin Alfa Solution for Injection(FSH) | Merck Serono | 928002 |
| Bovine Serum Albumin(BSA) | Sigma | A1933-25G |
| Fetuin | Sigma | F3004-1G |
| ITS | Sigma | 13146-5ML |
| Midkine human | Sigma | SRP3114 |
